# Differential effect of Fas activation on spinal muscular atrophy motoneuron death and induction of axonal growth

**DOI:** 10.1101/2022.05.12.491475

**Authors:** Salim Benlefki, Richard Younes, Désiré Challuau, Nathalie Bernard-Marissal, Cécile Hilaire, Frédérique Scamps, Melissa Bowerman, Rashmi Kothary, Bernard L Schneider, Cédric Raoul

## Abstract

Amyotrophic lateral sclerosis (ALS) and spinal muscular atrophy (SMA) are the most common motoneuron diseases affecting adults and infants, respectively. ALS and SMA are both characterized by the selective degeneration of motoneurons. Although different in their genetic etiology, growing evidence indicate that they share molecular and cellular pathogenic signatures that constitute potential common therapeutic targets. We previously described a motoneuron-specific death pathway elicited by the Fas death receptor, whereby vulnerable ALS motoneurons show an exacerbated sensitivity to Fas activation. However, the mechanisms that drive the loss of SMA motoneurons remains poorly understood. Here, we describe an *in vitro* model of SMA-associated degeneration using primary motoneurons derived from *Smn*^*2B/-*^ SMA mice and show that Fas activation selectively triggers death of the proximal motoneurons. Fas-induced death of SMA motoneurons has the molecular signature of the motoneuron-selective Fas death pathway that requires activation of p38 kinase, caspase-8, -9 and -3 as well as upregulation of collapsin response mediator protein 4 (CRMP4). In addition, Rho Kinase (ROCK) is required for Fas recruitment. Remarkably, we found that exogenous activation of Fas also promotes axonal elongation in both wildtype and SMA motoneurons. Axon outgrowth of motoneurons promoted by Fas requires the activity of ERK, ROCK and caspases. This work defines a dual role of Fas signaling in motoneurons that can elicit distinct responses from cell death to axonal growth.

## 1. INTRODUCTION

Motoneuron diseases (MNDs) constitute a group of devastating neurodegenerative disorders characterized by the progressive and selective degeneration of motoneurons in the brain and/or spinal cord. Amongst MNDs, spinal muscular atrophy (SMA) and amyotrophic lateral sclerosis (ALS) are the most common in children and adults, respectively. SMA is an inherited monogenic disease caused by loss-of-function mutations in the *survival motor neuron* (*SMN1*) gene and characterized by selective proximal muscle weakness and atrophy (Lefebvre *et al*., 1995; Crawford & Pardo, 1996). SMA is a clinically heterogeneous disease with various types (type 0-IV), which are classified according to age of onset and severity of symptoms. By contrast, ALS is inherited in approximately 10% of cases. With over 30 genes described to date, ALS causing mutations are most frequently found in *chromosome 9 open reading frame 72* (*C9ORF72*), *superoxide dismutase-1* (*SOD1*), *fused in sarcoma* (*FUS*) and *TAR DNA binding* protein (*TARDBP*) genes (Shatunov & Al-Chalabi, 2021). Familial cases are clinically indistinguishable from sporadic cases, both of which display upper and lower motoneuron signs as a common feature, which also illustrate the biological complexity of the disease.

As SMA and ALS are genetically distinct, they have traditionally been considered as separate disorders (Farrar & Kiernan, 2015; Shatunov & Al-Chalabi, 2021). However, accumulating evidence suggests that the convergence of aberrant pathways underpins the selective vulnerability of motoneurons in the context of ubiquitously expressed mutant proteins. Indeed, SMA and ALS share pathological features such as excitability defects in motoneurons that implicate intrinsic and extrinsic determinants, differential sprouting ability that underlie the selective vulnerability of motoneuron populations, non-cell-autonomous mechanisms where astrocytes and microglial cells produce neuroinflammatory and deleterious factors including tumor necrosis factor alpha, nitric oxide and interleukin-1β (Bowerman *et al*., 2017). More recently, a transcriptomic study conducted on vulnerable motoneurons isolated from the spinal cord of the *Smn*^*2B/-*^ mouse model of SMA (Bowerman *et al*., 2012a), has revealed a significant upregulation of the Fas (CD95) death receptor (Murray *et al*., 2015). Furthermore, motoneurons derived from SMA patient-induced pluripotent stem cells, which degenerate over time in culture, show upregulation of Fas ligand (FasL) and activation of downstream caspases (Sareen *et al*., 2012). Interestingly, we have previously demonstrated that Fas triggers a motoneuron-restricted death pathway that is exacerbated by ALS-linked mutant SOD1 (Raoul *et al*., 1999; Raoul *et al*., 2002; Raoul *et al*., 2006). The underlying pathway implicates the activation of p38 kinase and caspases as well as upregulation of collapsin response mediator protein 4 (Crmp4), a protein implicated in axonal degeneration (Duplan *et al*., 2010). Thus, the increased death activity of Fas in SMA and ALS could be a novel link between these two genetically distinct motoneuron diseases.

Here, we initially investigated whether Fas signaling was involved in the death of SMA motoneurons. We show that a population of primary motoneurons purified from *Smn*^*2B/-*^ mice progressively degenerate in culture. We observed the selective death of proximal *Smn*^*2B/-*^ motoneurons that can be blocked by interfering with the endogenous Fas-FasL interaction, activation of caspases, p38 kinase, Rho kinase (ROCK) and CRMP4 upregulation. Unexpectedly, we observed that application of exogenous soluble FasL also induces axonal growth in both wildtype and SMA motoneurons. Fas-dependent axon extension implicates the extracellular signal-related protein kinase (ERK) and caspases.

## 2. MATERIALS AND METHODS

### 2.1 Animals

All animal experiments were approved by the national ethics committee on animal experimentation and were performed in compliance with the European community and national directives for the care and use of laboratory animals. *Smn*^*2B/2B*^ and *Smn*^*+/-*^ mice were maintained on a C57BL/6J background under specific-pathogen-free conditions. *Smn*^*2B/2B*^ were bred with *Smn*^*+/-*^ mice to generate progeny with the *Smn*^*2B/+*^ healthy controls and *Smn*^*2B/-*^ SMA mice (Bowerman *et al*., 2012a).

### 2.2 Reagents

Soluble recombinant human FasL, enhancer for ligand and recombinant human Fas-Fc were purchased from Enzo Life Sciences. z-IETD-fmk, z-DEVD-fmk, Ac-LEHD-cmk, SB203580, PD98059 and Y-27632 were purchased from Merck. Rabbit polyclonal antibodies against Forkhead box P1 (FoxP1) (ab16645) and CRMP4 (ab101009) were purchased from Abcam. Monoclonal antibodies anti-Islet-1 and anti-Islet-2 (4D5 and 2D6) were from Developmental Studies Hybridoma Bank of Iowa University. Rabbit polyclonal antibodies against β-tubulin III (T2200) was from Sigma-Aldrich. Chicken polyclonal antibodies against GFP (ab13970) were purchased from Abcam.

### 2.3 Motoneuron culture

Mice at embryonic day (E)12.5 were kept on ice in Hibernate E (ThermoFisher Scientific) and the genotype were determined by PCR of tail DNA (Hammond *et al*., 2010). Motoneurons were isolated from the spinal cord of *Smn*^*2B/+*^ *and Smn*^*2B/-*^ embryos using 5.2% iodixanol density gradient centrifugation combined with p75-based magnetic cell isolation (Miltenyi Biotec) as we previously described (Soulard *et al*., 2020). Motoneurons from the spinal cords of *Hb9::GFP* embryos were isolated using iodixanol density gradient centrifugation. Motoneurons were plated on poly-ornithine/laminin-treated wells in the presence of a cocktail of neurotrophic factors (0.1 ng/ml glial-derived neurotrophic factor, 1 ng/ml brain-derived neurotrophic factor and 10 ng/ml ciliary neurotrophic factor in supplemented Neurobasal medium (ThermoFisher Scientific)). Supplemented Neurobasal medium contains 2% (vol/vol) horse serum, 25 mM L-glutamate, 25 mM β-mercaptoethanol, 0.5 mM L-glutamine, 2% (vol/vol) B-27 supplement (ThermoFisher Scientific) and 0.5% penicillin/streptomycin.

### 2.4 Immunocytochemistry

Motoneurons were plated on poly-ornithine/laminin-treated glass coverslips at the density of 5000 cells per cm^2^. Neurons were incubated for 15 min on ice in 2% paraformaldehyde (PFA) in phosphate-buffered saline (PBS)-culture medium (1:1) and for an additional 15 min in 4% PFA in PBS. Cells were washed with PBS and incubated for 1 h at room temperature in PBS containing 4% BSA, 4% donkey serum and 0.1% Triton X-100. Cells were incubated overnight at +4°C with the primary antibodies diluted in PBS containing 4% BSA, 4% donkey serum and 0.1% Triton X-100. After PBS washes, cells were incubated with fluorophore-conjugated secondary antibodies (ThermoFisher Scientific) for 1 h at room temperature. Cells were then washed in PBS and incubated for 5 min in PBS containing 4′,6-diamidino-2-phenylindole dihydrochloride (DAPI). Coverslips were mounted onto glass slides using Mowiol medium.

### 2.5 Production of recombinant adeno-associated virus

Adeno-associated-viral (AAV) vectors containing short hairpin RNA (shRNA) targeting Crmp4 or control mismatch were previously developed and validated (Duplan *et al*., 2010). The shRNA stem sequence used are: shCrmp4, 5’-GAACCTGAGTCTAGCCTGA-3’ and shCrmp4-mismatch, 5’-GAACCTGAGTATATCCTGA-3’. These shRNA were cloned into a bipartite pAAV vector with shRNA expression controlled by the H1 promoter and EGFP expression controlled by the cytomegalovirus promoter. AAV2/6 vectors were produced by the Bertarelli platform for Gene Therapy at EPFL (Lausanne, Switzerland). Briefly, pAAV plasmids were co-transfected with a pDP6 helper plasmid into HEK293-AAV cells (Agilent). Cells were lysed 72 h after transfection and viral particles were purified using Iodixanol gradient followed by separation ion-exchange chromatography (GE Healthcare). The infectivity titer of each virus (expressed as transduced units/ml) was determined following infection of HEK293T cells by real-time PCR using primers for beta-globin intron and human albumin.

### 2.6 Motoneuron survival and axon length

Motoneurons were seeded at a density of 750 cells/cm^2^. Three hours (0 DIV), 48 h (2 DIV) or 72 h (3 DIV) after plating, the number of surviving motoneurons were determined by counting two diameters. Between 60 and 150 motoneurons were counted in 2-cm diameters of each well by fluorescence or phase contrast microscopy using morphological criteria (Raoul *et al*., 2002). To compare values between different experiments, survival is expressed relative to the control condition where motoneurons are cultured only in the presence of the cocktail of neurotrophic factors. For the quantification of axon length, motoneurons were seeded at a density of 750 cells/cm^2^. Motoneurons were then processed for immunostaining with anti-GFP (for *Hb9::GFP* motoneurons) or β-tubulin III (for *Smn*^*2B/-*^ motoneurons) antibodies. Images were acquired with a ZEISS Axio Imager Z2 Apotome and axon length was determined using the ImageJ software and NeuronJ plugin (National Institutes of Health, USA)(Otsmane *et al*., 2014). Total axon length was determined by measuring the length of the longest neurite with connected branches of GFP-positive or β-tubulin III-positive motoneurons.

### 2.7 Statistical analyses

Statistical significance was determined by unpaired two-tailed *t* test, one-way or two-way analysis of variance (ANOVA) followed by a Tukey’s *post hoc* test. Statistical analyses were done with Graphpad Prism software (GraphPad Software). Significance was accepted at the level of *P* < 0.05.

## 3. RESULTS

### 3.1 Selective death of Smn-depleted proximal motoneurons *in vitro*

To study how Smn depletion impacts motoneuron death, we first isolated spinal motoneurons from *Smn*^*2B/+*^, *Smn*^*2B/-*^ and wildtype embryos and determined their survival over time in culture. While the survival of *Smn*^*2B/+*^ motoneurons is comparable to that of wildtype and remains stable over time, a notable decrease in the survival of *Smn*^*2B/-*^ motoneurons can be observed 2 and 3 days after seeding (Figure 1a). Thus, motoneurons from SMA mice show an altered survival *in vitro*, although the survival of about 60% of them is not affected by Smn depletion.

**FIGURE 1.**
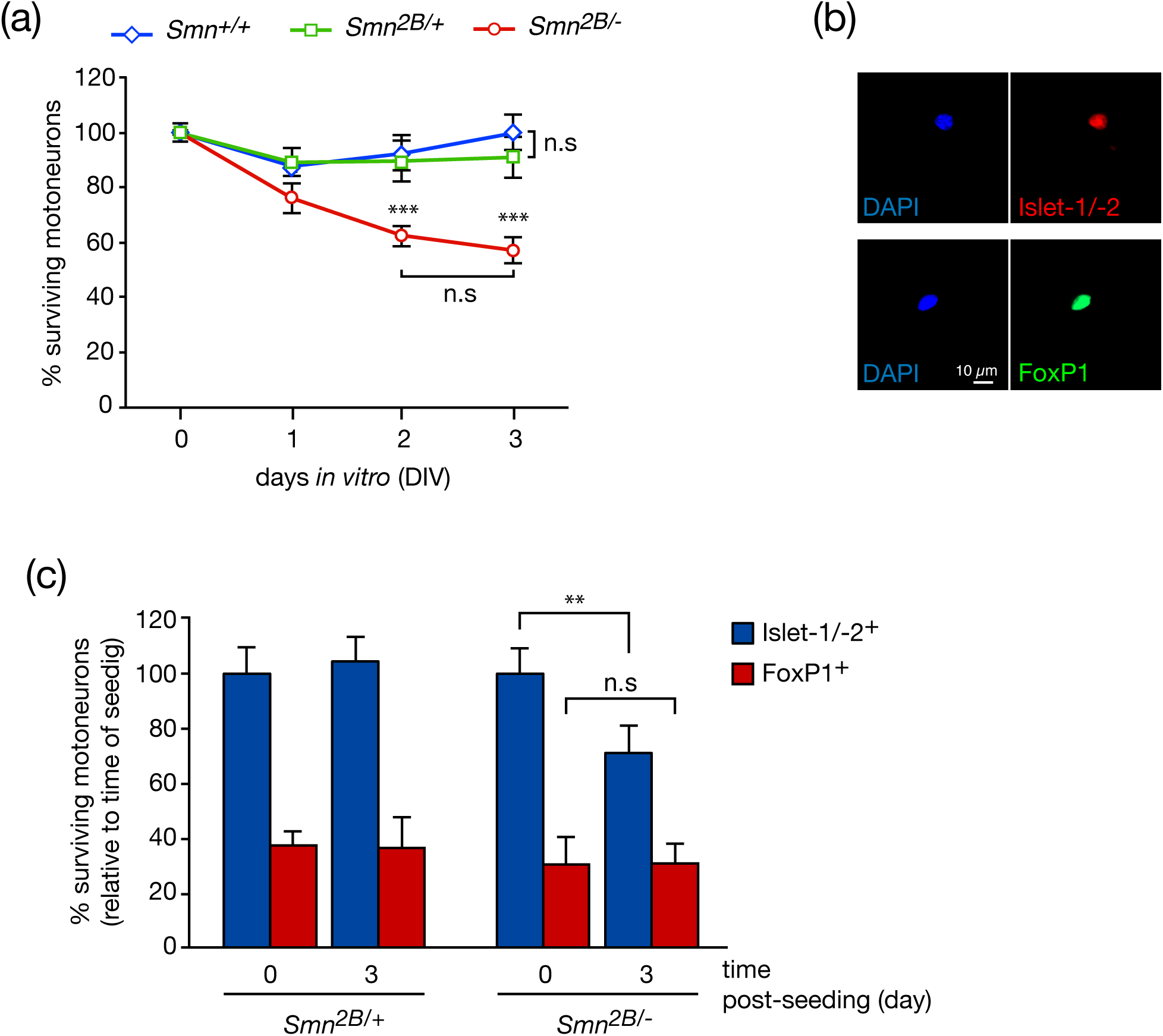
Selective death of *Smn*^*2B/-*^ motoneurons *in vitro*. **(a)** Survival of motoneurons immunopurified from wildtype (*Smn*^*+/+*^), *Smn*^*2B/+*^ and *Smn*^*2B/-*^ embryonic spinal cord. Motoneurons were cultured in the presence of neurotrophic factors and the percentage of surviving neurons was determined at the indicated time. For each genotype, the survival is expressed relative to the survival 3 hours after plating (0 DIV). **(b)** Illustrative immunostaining of *Smn*^*2B/-*^ motoneurons, cultured for 3 days, with anti-Islet-1, anti-Islet-2 or anti-FoxP1 antibodies. Scale bar, 10 μm. **(c)** *Smn*^*2B/+*^ and *Smn*^*2B/-*^ motoneurons were cultured for 0 and 3 DIV and immunostained with anti-Islet-1/-2 and FoxP1 antibodies. The proportion of FoxP1^+^ motoneurons is expressed as the percentage of the number of Islet-1/-2^+^ neurons at 0 DIV (time of seeding) for each genotype. Values are means ± standard error of the mean (SEM), \*\**P* < 0.01, \*\*\**P* < 0.001, n.s, non-significant, 1-way (c) or 2-way (a) ANOVA with Tukey’s *post hoc* test. Data are representative of 3-4 independent experiment, each done in triplicate. Statistical attributes are shown for *Smn*^*+/+*^ vs *Smn*^*2B/-*^. *Smn*^*2B/+*^ vs *Smn*^*2B/-*^ at 2 DIV, \*\**P* < 0.01 and *Smn*^*2B/-*^ vs *Smn*^*2B/+*^ at 3 DIV, \*\*\**P* < 0.001. *Smn*^*+/+*^ and *Smn*^*2B/+*^ are statistically indistinguishable at all times of culture.

Distal limb muscles are less affected than proximal muscles in SMA (Burr & Reddivari, 2021). FoxP1 transcription factor is selectively expressed by lateral motor column (LMC) neurons that innervate distal limb muscles (Dasen *et al*., 2008). We asked whether LMC motoneurons have a differential susceptibility to Smn depletion. We immunostained *Smn*^*2B/+*^ and *Smn*^*2B/-*^ motoneurons for FoxP1 and the LIM homeodomain transcription factors Islet-1 and Islet-2 that are expressed by all somatic motoneurons (Figure 1b). We then determined the relative proportion of LMC neurons after 3 days *in vitro* (DIV) (Figure 1c). We observed that the proportion of FoxP1^+^ motoneurons was unchanged between the time of seeding and 3 DIV in control *Smn*^*2B/+*^ mice. As expected, the survival of Islet-1/-2^+^ *Smn*^*2B/-*^ motoneurons was significantly decreased at 3 DIV compared to seeding time. However, the relative proportion of FoxP1^+^ *Smn*^*2B/-*^ motoneurons remains unchanged between the time of seeding and 3 DIV (Figure 1c), suggesting an increased vulnerability to Smn depletion among the population of Islet-1/-2^+^ FoxP1^-^ proximal motoneurons.

### 3.2 Death of SMA motoneurons involves the motoneuron-selective Fas death signaling

Since Fas is upregulated in vulnerable spinal motoneurons of *Smn*^*2B/-*^ mice and has been proposed to induce death of human motoneurons derived from SMA induced pluripotent stem cells (Sareen *et al*., 2012; Murray *et al*., 2015), we investigated whether Fas might trigger death of embryonic *Smn*^*2B/-*^ motoneurons. When we blocked endogenous Fas-FasL interaction by the Fas-Fc antagonist (Raoul *et al*., 1999), we prevented motoneuron loss induced by Smn depletion (Figure 2a). It is noteworthy, that when Fas was exogenously activated by the soluble recombinant form of FasL (sFasL), the proportion of *Smn*^*2B/-*^ motoneurons that die after 2 DIV was not increased (Figure 2a). These results indicate that Fas triggers death of motoneurons in the same proportion as Smn depletion. To evaluate whether the increased vulnerability of the FoxP1^-^ population of *Smn*^*2B/-*^ motoneurons is an idiosyncrasy of Smn loss (Figure 1c), we treated wildtype motoneurons with sFasL and determined the percentage of death induced by Fas in FoxP1^-^ and Foxp1^+^ motoneuron populations. We did not observe any different vulnerability to Fas activation between the distal and proximal populations of wildtype motoneurons (Figure 2b).

**FIGURE 2.**
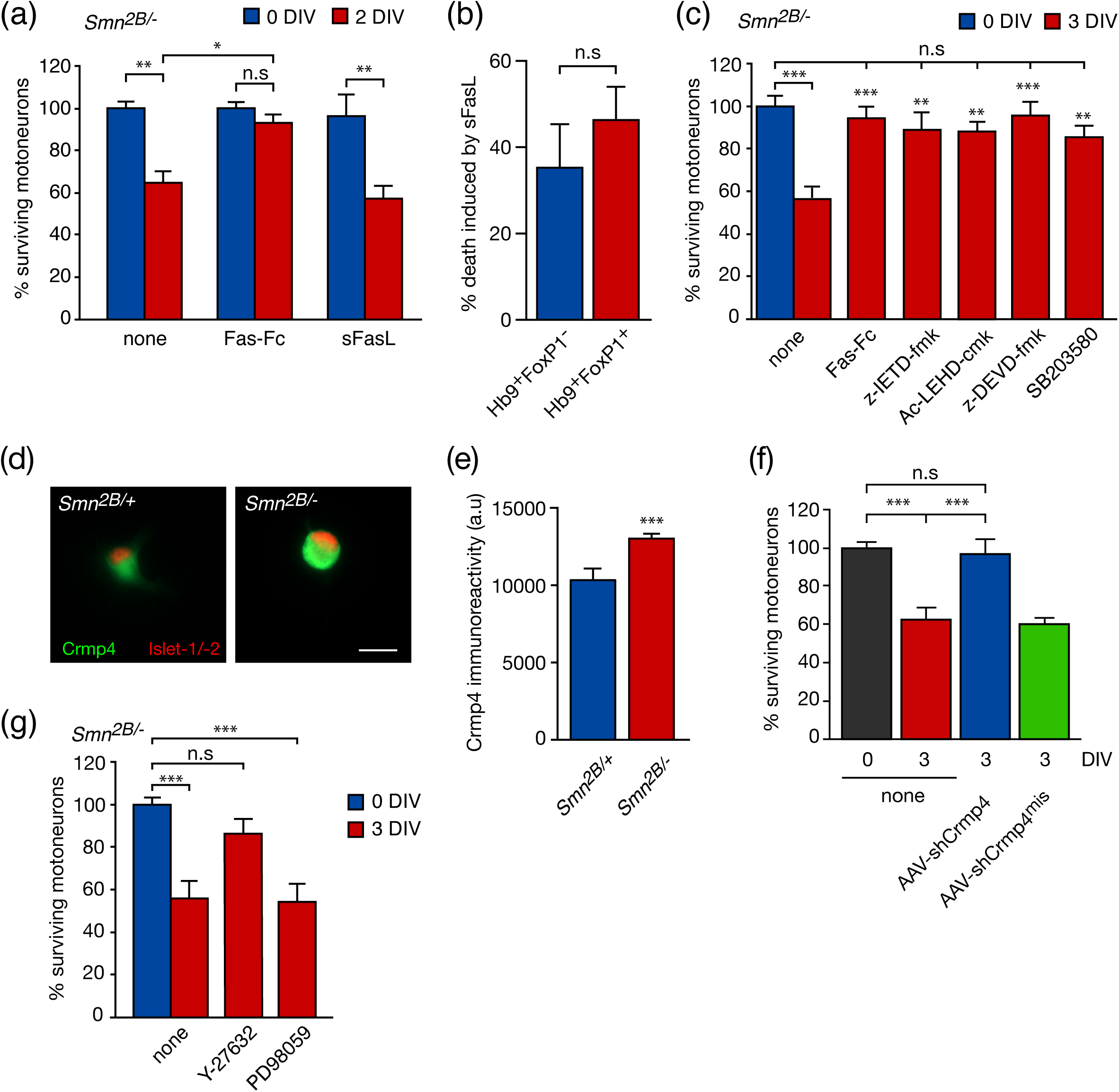
Death of *Smn*^*2B/-*^ motoneurons occurs through the Fas death pathway. **(a)** *Smn*^*2B/-*^ motoneurons were maintained in culture in the presence (or not) of Fas-Fc (1 μg/ml) or sFasL (100 ng/ml in the presence of 1 μg/ml enhancer antibody). Motoneuron survival was determined after 2 DIV and expressed relative to the number of motoneurons at the time of plating (0 DIV) in each condition. **(b)** Motoneurons from E12.5 spinal cord of *Hb9::GFP* embryos were cultured for 24 h and then treated (or not) with sFasL. The percentage of GFP^+^FoxP1^-^ and GFP^+^FoxP1^+^ motoneurons killed by Fas was determined 48 h later. **(c)** *Smn*^*2B/-*^ motoneurons were cultured in the presence of Fas-Fc (1 μg/ml), z-IETD-fmk (10 μM), Ac-LEHD-cmk (1 μM), z-DEVD-fmk (10 μM) or SB203580 (10 μM). Motoneuron survival was determined after 3 DIV and expressed relative to the number of motoneurons at the time of plating (0 DIV, none). **(d)** *Smn*^*2B/+*^ and *Smn*^*2B/-*^ motoneurons were cultured for 24 h and immunostained with anti-Islet-1/-2 and CRMP4 antibodies. Scale bar, 25 μm. **(e)** For the quantification of CRMP4 immunostaining, the CRMP4 mean fluorescence is expressed relative to Islet-1/-2 fluorescence (a.u, arbitrary unit). A total of 90 motoneurons were quantified by condition (*n* = 3). **(f)** SMA motoneurons were infected with 10 TU per cell of AAV-shCrmp4 and AAV-shCrmp4^mis^ after plating. The survival of motoneurons was determined at 3 DIV, expressed relative to the control condition (none) at the time of seeding (0 DIV). **(g)** *Smn*^*2B/-*^ motoneurons were incubated (or not) with 10 μM of Y-27632 and 10 μM of PD98059 for 3 days. Motoneuron survival was expressed relative to the control condition at 0 DIV. Values are means ± SEM, \**P* < 0.05, \*\**P* < 0.01, \*\*\**P* < 0.001, n.s, non-significant, ANOVA with Tukey’s *post hoc* test (a,c,f,g) or t test (b,e). Data are representative of at least three independent experiments, each done in triplicate.

We previously demonstrated that Fas-induced death of motoneurons involves the FADD-Caspase-8 pathway that synergistically acts with a Daxx-p38 kinase pathway (Raoul *et al*., 2002). To test whether Fas signaling involves the classical caspase-8, -9 and -3 caspases in *Smn*^*2B/-*^ motoneuron death, neurons were treated with their respective inhibitors. We observed that death of SMA motoneurons was completely blocked by the z-IETD-fmk caspase-8 inhibitor, Ac-LEHD-cmk caspase-9 inhibitor and the z-DEVD-fmk caspase-3 inhibitor (Figure 2c). We then inhibited p38 kinase activity using SB203580, which completely prevented Smn-dependent death of motoneurons (Figure 2c). Consistently, the protective effect of Fas-Fc on SMA motoneurons was also observed after 3 DIV. Another signature of the motoneuron-specific Fas signaling is the upregulation of CRMP4 that we showed to contribute to neuronal degeneration both *in vitro* and *in vivo* (Duplan *et al*., 2010). To investigate whether the death of SMA motoneurons involves CRMP4, we first determined CRMP4 levels in *Smn*^*2B/-*^ motoneurons. Quantification of fluorescence intensity show increased levels of CRMP4 in *Smn*^*2B/-*^ compared to *Smn*^*2B/+*^ motoneurons (Figure 2d,e). We then explored the functional implication of CRMP4 in the death of SMA motoneurons. We have previously shown that the silencing of CRMP4, through AAV6 vectors encoding an shRNA against *Crmp4 (*AAV*-*shCrmp4) that can transduce up to 90% of motoneurons *in vitro* (Langou *et al*., 2010), prevents death of motoneurons induced by the Fas pathway (Duplan *et al*., 2010). Reduction of CRMP4 expression in *Smn*^*2B/-*^ through AAV-shCrmp4 protected motoneurons from death induced by Smn depletion. Expression of a mismatch control (AAV-shCrmp4^mis^) had no effect on the survival of SMA motoneurons (Figure 2f). These results indicate that Smn depletion activates components of the motoneuron-restricted Fas death signaling.

The intracellular pathways underlying motoneuron degeneration under conditions of Smn depletion remain poorly understood. Combination of Fasudil that inhibits ROCK and selumetinib that inhibits ERK activity in SMA mouse neonates suggested that a ROCK to ERK cross-talk contribute to SMA pathogenesis (Hensel *et al*., 2017). We then asked whether ROCK and ERK pathway were implicated in the death of SMA primary motoneurons. While inhibition of the ROCK pathway by Y-27632 prevented death of motoneurons isolated from *Smn*^*2B/-*^ mice, inhibition of the ERK pathway by PD98059 had no effect on their survival (Figure 2g).

### 3.3 Exogenous Fas activation elicits axon outgrowth in motoneurons

The complexity of Fas signaling pathways is illustrated by its ability to promote neurite outgrowth in sensory neurons or increase the branching number in hippocampal neurons. These neuronal types are however resistant to Fas-mediated apoptosis (Desbarats *et al*., 2003; Zuliani *et al*., 2006). Because we previously demonstrated that activation of the lymphotoxin beta receptor (LT-βR) by its ligand LIGHT can elicit opposite responses in motoneurons by inducing either death or axon outgrowth (Otsmane *et al*., 2014), we examined the effect of Fas engagement by sFasL on axonal growth. The Fas-mediated death of motoneurons requires at least 2 DIV (Raoul *et al*., 1999). This observation is consistent with our finding that *Smn*^*2B/-*^ motoneuron death is not observed before 2 DIV (Figure 1a). Total axon length was thus determined following 24 h of treatment with sFasL. We observed that Fas activation significantly increases axon length in motoneurons (Figure 3a,b). We then asked whether this dual response to Fas stimulation could be impacted by Smn depletion. We found that exogenous Fas activation also promotes axonal elongation in *Smn*^*2B/-*^ motoneurons (Figure 3c).

**FIGURE 3.**
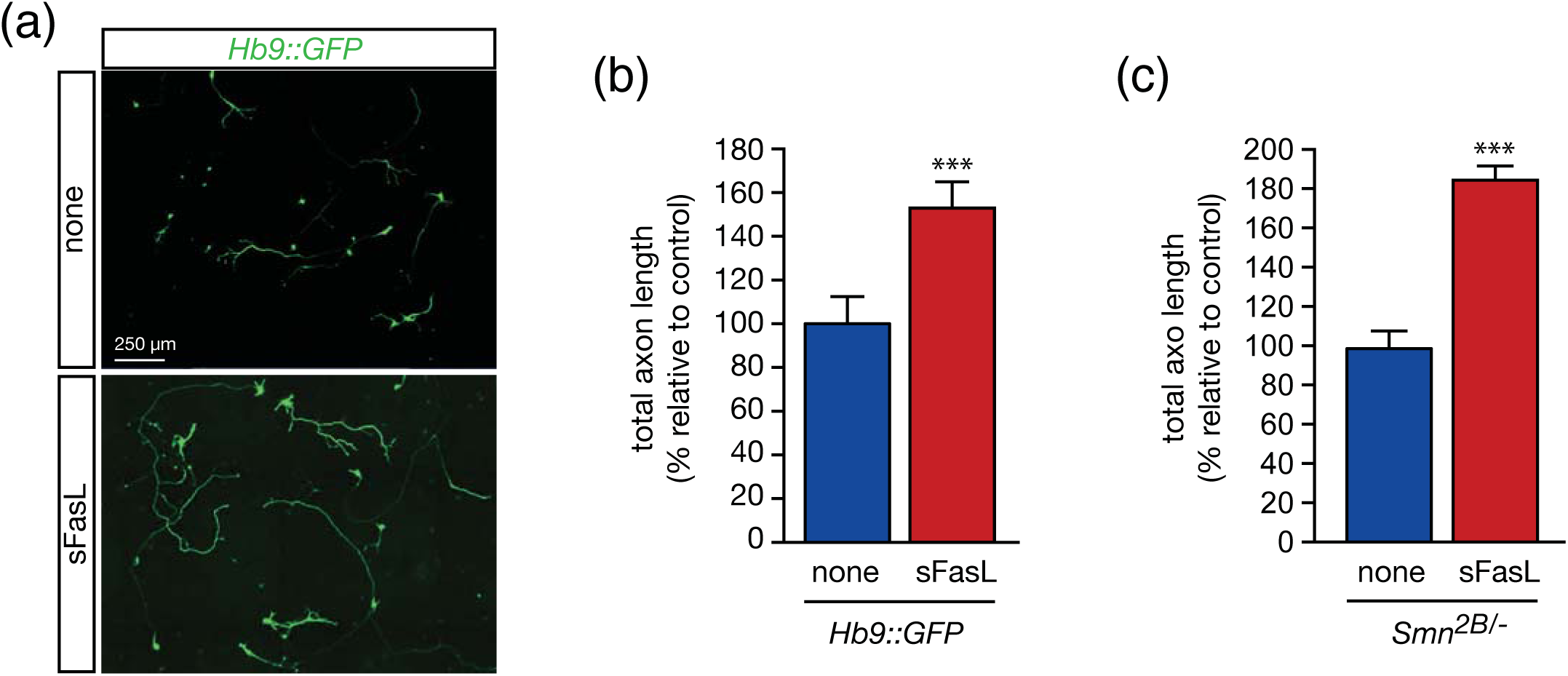
Fas activation induces axonal elongation in motoneurons. **(a)** Motoneurons were isolated from *Hb9::GFP* embryos and cultured for 24 h before being incubated with 100 μg/ml of sFasL (and 1 μg/ml enhancer antibody) for 24 h. Representative image; scale bar, 250 μm. **(b)** The total axon length, expressed relative to the control condition, was calculated after 24 h of treatment with sFasL. **(c)** The measurement of axon length was performed in *Smn*^*2B/-*^ motoneurons 24 h after treatment with sFasL. Data are represented as means ± SEM, experiences were repeated independently at least three time, each in triplicate, unpaired two-tailed *t* test, \*\*\**P* < 0.001.

### ERK and caspase signaling are required for Fas-induced axon outgrowth

We previously shown that LT-βR activation in motoneurons elicits the ERK signaling pathway required for neurite outgrowth, but not cell death (Otsmane *et al*., 2014). To evaluate the relative contribution of the ERK pathway in Fas-induced axon outgrowth in motoneurons, we pharmacologically inhibited ERK with PD98059 following sFasL treatment. This reduced the total axon length of motoneurons following Fas activation (Figure 4a). However, pharmacological inhibition of the ERK pathway had no effect on Fas-dependent death of motoneurons (Figure 4b). The serine/threonine kinase ROCK regulates actin cytoskeleton dynamics and prevents axonal growth (Tonges *et al*., 2011).

**FIGURE 4.**
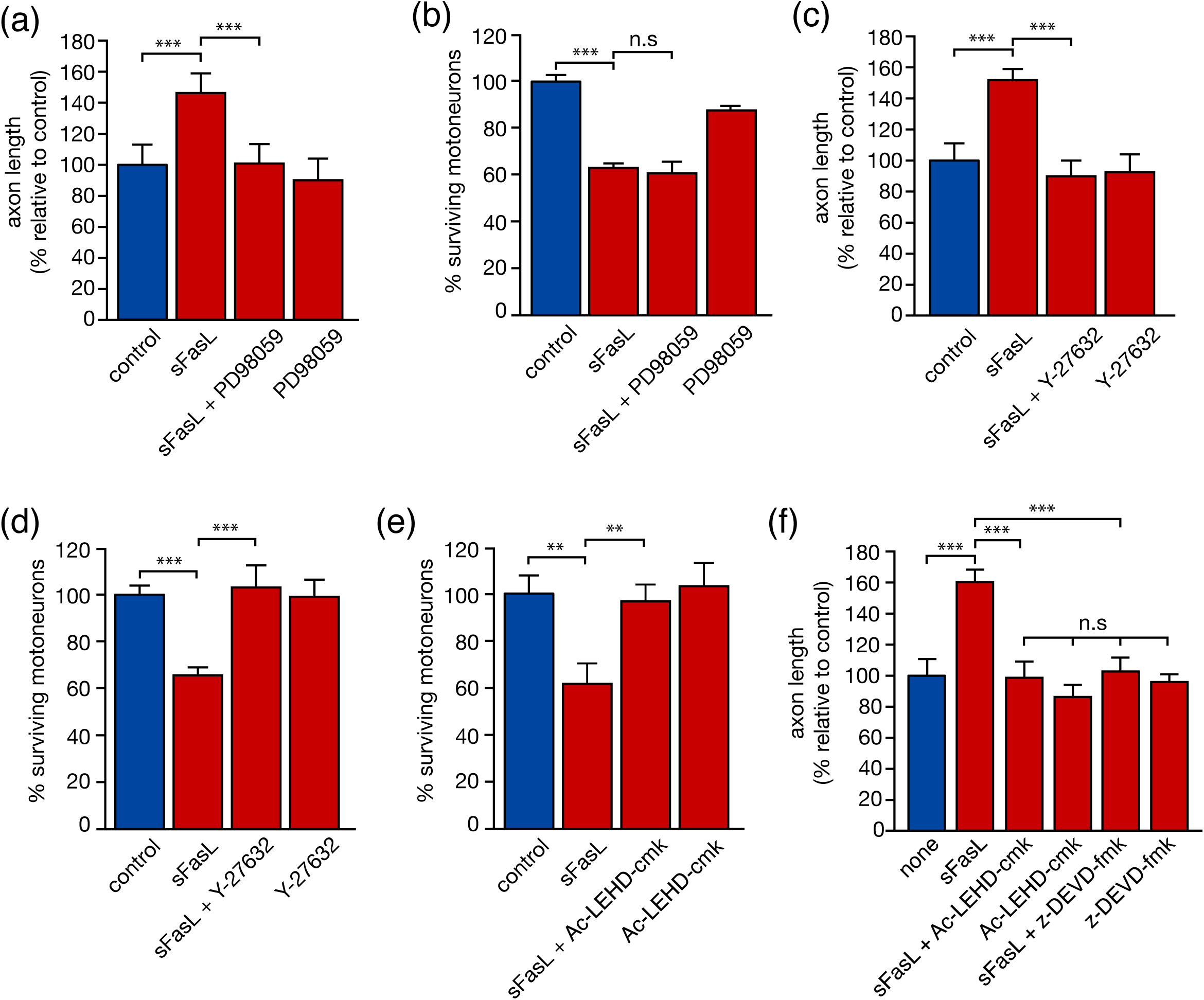
Fas-induced axon outgrowth requires ERK and Caspase activity. **(a)** *Hb9::GFP* motoneurons were maintained in culture for 24 h and treated with sFasL (100 ng/ml and 1 μg/ml enhancer antibody) in combination with PD98059 (10 μM). Total axon length was calculated 24 h later. **(b)** Motoneurons were cultured for 24 h and treated with sFasL and PD98059. Surviving GFP^+^ motoneurons were directly counted under fluorescence microscope 48 h later. **(c**,**d)** Twenty-four hours post seeding, sFasL (100 ng/ml) and Y-27632 (10 μM) were added to *Hb9::GFP* motoneurons. **(c)** Total axon length was determined 24 h later. **(d)** Motoneuron survival was determined 48 h later. **(e)** Motoneurons were maintained in culture for 24 h and treated with Fas activator and caspase-9 inhibitor Ac-LEHD-cmk (1 μM). Motoneuron survival was determined 48 h later and expressed relative to the non-treated condition. **(f)** Motoneurons were treated (or not) with sFasL in combination with Ac-LEHD-cmk (1 μM) or z-DEVD-fmk (10 μM). Axon length was determined 24 h later. The graphs show the mean values ± SEM of at least three independent experiments performed in triplicate, \*\**P* < 0.01, \*\*\**P* < 0.001, n.s, non-significant, ANOVA with Tukey’s *post hoc* test.

As shown above, death of *Smn*^*2B/-*^ motoneurons involves ROCK signaling (Figure 2g). We therefore asked whether ROCK inhibition might impact axon length as well as death of wildtype motoneurons following exogenous Fas activation. Addition of Y-27632 to motoneuron cultures treated with sFasL abolished Fas-induced axonal elongation (Figure 4c), as well as Fas-induced death of motoneurons (Figure 4d). This observation is consistent with the role of ROCK in Fas clustering (see Discussion).

Accumulating evidence shows that the function of executioner caspases is not limited to apoptotic activity but may also be involved in neurite growth as well as axonal guidance (Hollville & Deshmukh, 2018). We have previously shown that z-DEVD-fmk caspase-3 inhibitor saves motoneuron from Fas-induced death (Raoul *et al*., 1999). We also observed that Ac-LEHD-cmk caspase-9 inhibitor saves motoneuron from Fas-induced death (Figure 4e). We then applied Ac-LEHD-cmk and z-DEVD-fmk to assess the contribution of executioner caspase in Fas-induced neurite outgrowth. Total axonal length of motoneurons was significantly reduced when inhibitor of caspase-9 and caspase-3 were added with sFasL (Figure 4f). Combined, our results suggest that both the ERK and ROCK signaling pathways modulate the effect of Fas on axonal outgrowth but that only the latter contribute to Fas-dependent death of motoneurons. Caspases also seem to contribute to Fas-mediated axonal growth.

## 4. DISCUSSION

The molecular mechanisms that drive motoneuron death in SMA remain elusive, and those, like p53 pathway, which have been proposed as important are still the subject of conflicting results and debate (Simon *et al*., 2017; Courtney *et al*., 2019; Reedich *et al*., 2021). To explore SMA-related death mechanisms, we used the *Smn*^*2B/-*^ mouse model of SMA. *Smn*^*2B/-*^ mice harbor on one allele a three-nucleotide substitution mutation (termed 2B) within the exon splicing enhancer of *Smn* exon 7 and a null allele of *Smn* (Hammond *et al*., 2010). *Smn*^*2B/-*^ mice show reduced Smn levels in the brain and spinal cord to approximately 20% of control levels (Groen *et al*., 2018), and displays the pathological hallmarks of SMA observed in patients such as muscle weakness and gait abnormalities. These SMA mice have a median lifespan of 30 days and are characterized by motoneuron loss and neuromuscular defects (Bowerman *et al*., 2012a). We have shown here that motoneurons derived from *Smn*^*2B/-*^ mouse embryos have a survival that is progressively reduced with time in culture. This loss of motoneurons is dependent on the Fas-FasL interaction and preferentially involves the subpopulation that innervates proximal muscles, based on FoxP1 expression. This selective vulnerability of FoxP1^-^ proximal SMA motoneurons is consistent with the selective proximal distribution of muscle weakness observed in SMA patients and mice (Murray *et al*., 2008; Burr & Reddivari, 2021). The absence of FoxP1 expression, which we have used here for its role in defining columnar identity of motoneurons (Dasen *et al*., 2008), is also intriguing for other functional aspects related to a potential vulnerability to death. Indeed, levels of FoxP1 are decreased in the striatum of mice and patients with Huntington’s disease, and the forced expression of FoxP1 rescued cortical neurons from mutant Huntingtin-mediated toxicity (Louis Sam Titus *et al*., 2017). FoxP1 overexpression represses expression of proapoptotic genes in B cells (van Keimpema *et al*., 2014). FasL was identified as one of these proapoptotic genes directly repressed by FoxP1 (Dekker *et al*., 2019). Transcriptome analysis of a FoxP1-depleted human colon carcinoma cell line showed an upregulation of Forkhead box transcription factor class O (FOXO) target genes that include Fas (van Boxtel *et al*., 2013). If we refocus on the spinal motor system, FasL is a downstream target of Foxo3a to drive motoneuron death through Fas activation following neurotrophic factor deprivation (Barthelemy *et al*., 2004). Here, we observed that wildtype FoxP1^-^ and FoxP1^+^ motoneuron display similar susceptibly to Fas-induced death. Identity determinants, by engaging essential factors for neuronal diversification, projection and connectivity patterns, may also contribute, alongside interactions with genes such as Smn, to provide a cell-intrinsic program of vulnerability.

Death of Smn-depleted motoneurons can be prevented by ROCK, but not ERK, inhibitor. Studies on the contribution of ROCK in SMA mice have led to diverse conclusions (Bowerman *et al*., 2012b; Branchu *et al*., 2013; Hensel *et al*., 2017). This is likely related to the multi-systemic nature of the disease where various cell types contribute to the symptoms and may be the target of the inhibitors (Coque *et al*., 2014). The cellular modalities of ROCK and ERK inhibition (as well as culture conditions) may also help to explain the differences in ERK inhibitor effects between our observations made on immunopurified motoneurons and those made on mixed cultures of spinal cord obtained from the severe Taiwanese mice (Hensel *et al*., 2017). On the contribution of the ROCK pathway in Fas-dependent death of SMA motoneurons, our observation is reminiscent of what was observed in Jurkat cells where ROCK inhibition by Y-27632 reduced apoptosis induced by Fas stimulation or what was reported in human dermal microvascular endothelial cells where Fasudil, another ROCK inhibitor, protected from Fas-induced death (Arita *et al*., 2009). In the Jurkat cells, it was shown that upon Fas activation ROCK phosphorylates ezrin and meosin, thus linking Fas to the actin cytoskeleton and facilitating Fas clustering and formation of the death-inducing signaling complex (Hebert *et al*., 2008). Another evidence shows that the clustering of Fas, which occurs concomitantly with F-actin polarization and independently of the apoptotic program, requires ROCK activity (Soderstrom *et al*., 2005). Even more interesting is that Fas activation in Jurkat cells induces RhoA which acts upstream of ROCK. Our data support the growing interest of ROCK as a therapeutic target for motoneuron diseases, including ALS, in which we and others have shown the role of Fas (Aebischer *et al*., 2013). Of note, Fasudil is already being evaluated in a double-blind, randomized, placebo-controlled phase IIa trial in patients with ALS (Lingor *et al*., 2019). However, by acting very upstream in Fas signaling and abrogating therefore both deleterious (apoptosis) but also beneficial (axonal outgrowth) processes, the outcome of ROCK inhibition *in vivo* might also be interpreted under the light of Fas signaling.

Here, we report that Fas activation by the soluble form of FasL ligand can also elicit axon outgrowth in both wildtype and *Smn*^*2B/-*^ motoneurons. Previously, Fas stimulation was shown to increase the number of branching points, but not neurite length, of hippocampal neurons, independently of caspase activation (Zuliani *et al*., 2006). Another study also reported that Fas activation induced neurite outgrowth in sensory neurons in a caspase-8-independent, ERK pathway-dependent manner (Desbarats *et al*., 2003). We here reproduce this feature of dual signaling of a death receptor in motoneurons by highlighting that Fas requires ERK and caspases to elicit axon outgrowth, while it requires caspases, but not ERK, to trigger death. While the ERK pathway, which can modulate the local translation of proteins, is widely recognized as a key promoter of neurite outgrowth (Hausott & Klimaschewski, 2019), and localized caspase activity has been well documented during axon degeneration (Geden *et al*., 2019), the contribution of caspases to axonal elongation remains however elusive. The diversity of substrates of caspases already underlines the extent of their nonapoptotic function. Indeed, very recently, N-Terminomics has identified 906 caspase-3 and 124 caspase-9 substrates in healthy Jurkat cells (Araya *et al*., 2021). The majority were previously identified as apoptotic substrates, but other substrates were implicated in different pathways such as mRNA splicing, RNA metabolism, Notch-HLH pathway SUMOylation, HIV infection for caspase-3 and mRNA splicing, RNA metabolism, mitotic prometaphase, Rho GTPase signaling, and membrane trafficking for caspase-9. Another study in Jurkat cells following TRAIL activation, identified as caspase-3, -7, -8 substrates, proteins involved in endocytosis and vesicle trafficking (Agard *et al*., 2012). Additional caspase-3 substrate databases (MEROPS (Rawlings *et al*., 2010), CASBAH (Luthi & Martin, 2007), and DegraBase (Crawford *et al*., 2013)) have been described that illustrate the complexity of apoptotic and nonapoptotic caspase signaling. In *Xenopus*, it was shown that the chemotropic cue netrin-1 that induces retinal growth cone attraction, requires caspase-3 activity which is downstream of p38 kinase (Campbell & Holt, 2003). In chick embryos, caspase-3 is involved in the arborization pattern of developing axons of the ciliary ganglion (Katow *et al*., 2017). In rat, growth cone formation following axotomy of sensory and retinal axons implicates p38 kinase and caspase-3 activity (Verma *et al*., 2005). In mouse hippocampal neurons, the clustering of the neural cell adhesion molecule at the growth cone induces caspase-8 and -3 activation, which is required for neurite outgrowth likely through local proteolysis of the spectrin meshwork (Westphal *et al*., 2010). Thus, it will be imperative for future studies to examine the intracellular cascade that coordinates local protein translation, membrane remodeling and cytoskeleton dynamics in response to death receptor activation in motoneurons.

Our study reveals a common death mechanism that underpins the vulnerability of motoneurons to ALS-and SMA-causing genetic determinants. The intracellular relays of the motoneuron-restricted Fas death pathway are also recruited to execute death of SMA motoneurons. These common signaling intermediates may therefore appear as therapeutic targets for SMA. However, the duality of motoneuron response to Fas activation provides an important insight into the therapeutic approaches to be considered.

## Acknowledgments

We thank all members of the team, Dr. Lyndsay M Murray (University of Edinburgh) and Dr. Stéphane Nedelec (IFM, Paris) for their helpful comments throughout the work; Céline Salsac and Ariane Beauvais for their technical help and advices; the personnel of the Montpellier RIO imaging platform and the RAM-Neuro INM animal facility for their services.

## AUTHOR CONTRIBUTIONS

SB, MB and CR conceived and designed the analysis; SB and CR collected the data; SB, RY, DC, NBM, CR performed the experiments; RK and BS provided *Smn* mice and viral vectors respectively; SB, MB, RK, CH, FS, BS and CR analyzed the results; CR administrated the project, CR and SB drafted the manuscript and all authors participated in revisions. All authors reviewed the results and approved the final version of the manuscript.

## REFERENCES

Aebischer, J., Bernard-Marissal, N., Pettmann, B. & Raoul, C. (2013) Death Receptors in the Selective Degeneration of Motoneurons in Amyotrophic Lateral Sclerosis. J Neurodegener Dis, 2013, 746845.

Agard, N.J., Mahrus, S., Trinidad, J.C., Lynn, A., Burlingame, A.L. & Wells, J.A. (2012) Global kinetic analysis of proteolysis via quantitative targeted proteomics. Proc Natl Acad Sci U S A, 109, 1913–1918.

Araya, L.E., Soni, I.V., Hardy, J.A. & Julien, O. (2021) Deorphanizing Caspase-3 and Caspase-9 Substrates In and Out of Apoptosis with Deep Substrate Profiling. ACS Chem Biol.

Arita, R., Hata, Y., Nakao, S., Kita, T., Miura, M., Kawahara, S., Zandi, S., Almulki, L., Tayyari, F., Shimokawa, H., Hafezi-Moghadam, A. & Ishibashi, T. (2009) Rho kinase inhibition by fasudil ameliorates diabetes-induced microvascular damage. Diabetes, 58, 215–226.

Barthelemy, C., Henderson, C.E. & Pettmann, B. (2004) Foxo3a induces motoneuron death through the Fas pathway in cooperation with JNK. BMC Neurosci, 5, 48.

Bowerman, M., Murray, L.M., Beauvais, A., Pinheiro, B. & Kothary, R. (2012a) A critical smn threshold in mice dictates onset of an intermediate spinal muscular atrophy phenotype associated with a distinct neuromuscular junction pathology. Neuromuscul Disord, 22, 263–276.

Bowerman, M., Murray, L.M., Boyer, J.G., Anderson, C.L. & Kothary, R. (2012b) Fasudil improves survival and promotes skeletal muscle development in a mouse model of spinal muscular atrophy. BMC Med, 10, 24.

Bowerman, M., Murray, L.M., Scamps, F., Schneider, B.L., Kothary, R. & Raoul, C. (2017) Pathogenic commonalities between spinal muscular atrophy and amyotrophic lateral sclerosis: Converging roads to therapeutic development. Eur J Med Genet.

Branchu, J., Biondi, O., Chali, F., Collin, T., Leroy, F., Mamchaoui, K., Makoukji, J., Pariset, C., Lopes, P., Massaad, C., Chanoine, C. & Charbonnier, F. (2013) Shift from extracellular signal-regulated kinase to AKT/cAMP response element-binding protein pathway increases survival-motor-neuron expression in spinal-muscular-atrophy-like mice and patient cells. J Neurosci, 33, 4280–4294.

Burr, P. & Reddivari, A.K.R. (2021) Spinal Muscle Atrophy StatPearls, Treasure Island (FL).

Campbell, D.S. & Holt, C.E. (2003) Apoptotic pathway and MAPKs differentially regulate chemotropic responses of retinal growth cones. Neuron, 37, 939–952.

Coque, E., Raoul, C. & Bowerman, M. (2014) ROCK inhibition as a therapy for spinal muscular atrophy: understanding the repercussions on multiple cellular targets. Front Neurosci, 8, 271.

Courtney, N.L., Mole, A.J., Thomson, A.K. & Murray, L.M. (2019) Reduced P53 levels ameliorate neuromuscular junction loss without affecting motor neuron pathology in a mouse model of spinal muscular atrophy. Cell Death Dis, 10, 515.

Crawford, E.D., Seaman, J.E., Agard, N., Hsu, G.W., Julien, O., Mahrus, S., Nguyen, H., Shimbo, K., Yoshihara, H.A., Zhuang, M., Chalkley, R.J. & Wells, J.A. (2013) The DegraBase: a database of proteolysis in healthy and apoptotic human cells. Mol Cell Proteomics, 12, 813–824.

Crawford, T.O. & Pardo, C.A. (1996) The neurobiology of childhood spinal muscular atrophy. Neurobiol Dis, 3, 97–110.

Dasen, J.S., De Camilli, A., Wang, B., Tucker, P.W. & Jessell, T.M. (2008) Hox repertoires for motor neuron diversity and connectivity gated by a single accessory factor, FoxP1. Cell, 134, 304–316.

Dekker, J.D., Baracho, G.V., Zhu, Z., Ippolito, G.C., Schmitz, R.J., Rickert, R.C. & Tucker, H.O. (2019) Loss of the FOXP1 Transcription Factor Leads to Deregulation of B Lymphocyte Development and Function at Multiple Stages. Immunohorizons, 3, 447–462.

Desbarats, J., Birge, R.B., Mimouni-Rongy, M., Weinstein, D.E., Palerme, J.S. & Newell, M.K. (2003) Fas engagement induces neurite growth through ERK activation and p35 upregulation. Nat Cell Biol, 5, 118–125.

Duplan, L., Bernard, N., Casseron, W., Dudley, K., Thouvenot, E., Honnorat, J., Rogemond, V., De Bovis, B., Aebischer, P., Marin, P., Raoul, C., Henderson, C.E. & Pettmann, B. (2010) Collapsin response mediator protein 4a (CRMP4a) is upregulated in motoneurons of mutant SOD1 mice and can trigger motoneuron axonal degeneration and cell death. J Neurosci, 30, 785–796.

Farrar, M.A. & Kiernan, M.C. (2015) The Genetics of Spinal Muscular Atrophy: Progress and Challenges. Neurotherapeutics, 12, 290–302.

Geden, M.J., Romero, S.E. & Deshmukh, M. (2019) Apoptosis versus axon pruning: Molecular intersection of two distinct pathways for axon degeneration. Neurosci Res, 139, 3–8.

Groen, E.J.N., Perenthaler, E., Courtney, N.L., Jordan, C.Y., Shorrock, H.K., van der Hoorn, D., Huang, Y.T., Murray, L.M., Viero, G. & Gillingwater, T.H. (2018) Temporal and tissue-specific variability of SMN protein levels in mouse models of spinal muscular atrophy. Hum Mol Genet, 27, 2851–2862.

Hammond, S.M., Gogliotti, R.G., Rao, V., Beauvais, A., Kothary, R. & Didonato, C.J. (2010) Mouse Survival Motor Neuron Alleles That Mimic SMN2 Splicing and Are Inducible Rescue Embryonic Lethality Early in Development but Not Late. PLoS One, 5, e15887.

Hausott, B. & Klimaschewski, L. (2019) Promotion of Peripheral Nerve Regeneration by Stimulation of the Extracellular Signal-Regulated Kinase (ERK) Pathway. Anat Rec (Hoboken), 302, 1261–1267.

Hebert, M., Potin, S., Sebbagh, M., Bertoglio, J., Breard, J. & Hamelin, J. (2008) Rho-ROCK-dependent ezrin-radixin-moesin phosphorylation regulates Fas-mediated apoptosis in Jurkat cells. J Immunol, 181, 5963–5973.

Hensel, N., Baskal, S., Walter, L.M., Brinkmann, H., Gernert, M. & Claus, P. (2017) ERK and ROCK functionally interact in a signaling network that is compensationally upregulated in Spinal Muscular Atrophy. Neurobiol Dis, 108, 352–361.

Hollville, E. & Deshmukh, M. (2018) Physiological functions of non-apoptotic caspase activity in the nervous system. Semin Cell Dev Biol, 82, 127–136.

Katow, H., Kanaya, T., Ogawa, T., Egawa, R. & Yawo, H. (2017) Regulation of axon arborization pattern in the developing chick ciliary ganglion: Possible involvement of caspase 3. Dev Growth Differ, 59, 115–128.

Langou, K., Moumen, A., Pellegrino, C., Aebischer, J., Medina, I., Aebischer, P. & Raoul, C. (2010) AAV-mediated expression of wild-type and ALS-linked mutant VAPB selectively triggers death of motoneurons through a Ca2+-dependent ER-associated pathway. J Neurochem, 114, 795–809.

Lefebvre, S., Burglen, L., Reboullet, S., Clermont, O., Burlet, P., Viollet, L., Benichou, B., Cruaud, C., Millasseau, P., Zeviani, M. & et al. (1995) Identification and characterization of a spinal muscular atrophy-determining gene. Cell, 80, 155–165.

Lingor, P., Weber, M., Camu, W., Friede, T., Hilgers, R., Leha, A., Neuwirth, C., Gunther, R., Benatar, M., Kuzma-Kozakiewicz, M., Bidner, H., Blankenstein, C., Frontini, R., Ludolph, A., Koch, J.C. & Investigators, R.-A. (2019) ROCK-ALS: Protocol for a Randomized, Placebo-Controlled, Double-Blind Phase IIa Trial of Safety, Tolerability and Efficacy of the Rho Kinase (ROCK) Inhibitor Fasudil in Amyotrophic Lateral Sclerosis. Front Neurol, 10, 293.

Louis Sam Titus, A.S.C., Yusuff, T., Cassar, M., Thomas, E., Kretzschmar, D. & D’Mello, S.R. (2017) Reduced Expression of Foxp1 as a Contributing Factor in Huntington’s Disease. J Neurosci, 37, 6575–6587.

Luthi, A.U. & Martin, S.J. (2007) The CASBAH: a searchable database of caspase substrates. Cell death and differentiation, 14, 641–650.

Murray, L.M., Beauvais, A., Gibeault, S., Courtney, N.L. & Kothary, R. (2015) Transcriptional profiling of differentially vulnerable motor neurons at pre-symptomatic stage in the Smn (2b/-) mouse model of spinal muscular atrophy. Acta Neuropathol Commun, 3, 55.

Murray, L.M., Comley, L.H., Thomson, D., Parkinson, N., Talbot, K. & Gillingwater, T.H. (2008) Selective vulnerability of motor neurons and dissociation of pre-and post-synaptic pathology at the neuromuscular junction in mouse models of spinal muscular atrophy. Hum Mol Genet, 17, 949–962.

Otsmane, B., Moumen, A., Aebischer, J., Coque, E., Sar, C., Sunyach, C., Salsac, C., Valmier, J., Salinas, S., Bowerman, M. & Raoul, C. (2014) Somatic and axonal LIGHT signaling elicit degenerative and regenerative responses in motoneurons, respectively. EMBO Rep, 15, 540–547.

Raoul, C., Buhler, E., Sadeghi, C., Jacquier, A., Aebischer, P., Pettmann, B., Henderson, C.E. & Haase, G. (2006) Chronic activation in presymptomatic amyotrophic lateral sclerosis (ALS) mice of a feedback loop involving Fas, Daxx, and FasL. Proc Natl Acad Sci U S A, 103, 6007–6012.

Raoul, C., Estevez, A.G., Nishimune, H., Cleveland, D.W., deLapeyriere, O., Henderson, C.E., Haase, G. & Pettmann, B. (2002) Motoneuron death triggered by a specific pathway downstream of Fas. Potentiation by ALS-linked SOD1 mutations. Neuron, 35, 1067–1083.

Raoul, C., Henderson, C.E. & Pettmann, B. (1999) Programmed cell death of embryonic motoneurons triggered through the Fas death receptor. J Cell Biol, 147, 1049–1062.

Rawlings, N.D., Barrett, A.J. & Bateman, A. (2010) MEROPS: the peptidase database. Nucleic Acids Res, 38, D227–233.

Reedich, E.J., Kalski, M., Armijo, N., Cox, G.A. & DiDonato, C.J. (2021) Spinal motor neuron loss occurs through a p53-and-p21-independent mechanism in the Smn(2B/-) mouse model of spinal muscular atrophy. Exp Neurol, 337, 113587.

Sareen, D., Ebert, A.D., Heins, B.M., McGivern, J.V., Ornelas, L. & Svendsen, C.N. (2012) Inhibition of apoptosis blocks human motor neuron cell death in a stem cell model of spinal muscular atrophy. PLoS One, 7, e39113.

Shatunov, A. & Al-Chalabi, A. (2021) The genetic architecture of ALS. Neurobiol Dis, 147, 105156.

Simon, C.M., Dai, Y., Van Alstyne, M., Koutsioumpa, C., Pagiazitis, J.G., Chalif, J.I., Wang, X., Rabinowitz, J.E., Henderson, C.E., Pellizzoni, L. & Mentis, G.Z. (2017) Converging Mechanisms of p53 Activation Drive Motor Neuron Degeneration in Spinal Muscular Atrophy. Cell Rep, 21, 3767–3780.

Soderstrom, T.S., Nyberg, S.D. & Eriksson, J.E. (2005) CD95 capping is ROCK-dependent and dispensable for apoptosis. J Cell Sci, 118, 2211–2223.

Soulard, C., Salsac, C., Mouzat, K., Hilaire, C., Roussel, J., Mezghrani, A., Lumbroso, S., Raoul, C. & Scamps, F. (2020) Spinal Motoneuron TMEM16F Acts at C-boutons to Modulate Motor Resistance and Contributes to ALS Pathogenesis. Cell Rep, 30, 2581–2593 e2587.

Tonges, L., Koch, J.C., Bahr, M. & Lingor, P. (2011) ROCKing Regeneration: Rho Kinase Inhibition as Molecular Target for Neurorestoration. Front Mol Neurosci, 4, 39.

van Boxtel, R., Gomez-Puerto, C., Mokry, M., Eijkelenboom, A., van der Vos, K.E., Nieuwenhuis, E.E., Burgering, B.M., Lam, E.W. & Coffer, P.J. (2013) FOXP1 acts through a negative feedback loop to suppress FOXO-induced apoptosis. Cell death and differentiation, 20, 1219–1229.

van Keimpema, M., Gruneberg, L.J., Mokry, M., van Boxtel, R., Koster, J., Coffer, P.J., Pals, S.T. & Spaargaren, M. (2014) FOXP1 directly represses transcription of proapoptotic genes and cooperates with NF-kappaB to promote survival of human B cells. Blood, 124, 3431–3440.

Verma, P., Chierzi, S., Codd, A.M., Campbell, D.S., Meyer, R.L., Holt, C.E. & Fawcett, J.W. (2005) Axonal protein synthesis and degradation are necessary for efficient growth cone regeneration. J Neurosci, 25, 331–342.

Westphal, D., Sytnyk, V., Schachner, M. & Leshchyns’ka, I. (2010) Clustering of the neural cell adhesion molecule (NCAM) at the neuronal cell surface induces caspase-8-and -3-dependent changes of the spectrin meshwork required for NCAM-mediated neurite outgrowth. J Biol Chem, 285, 42046–42057.

Zuliani, C., Kleber, S., Klussmann, S., Wenger, T., Kenzelmann, M., Schreglmann, N., Martinez, A., del Rio, J.A., Soriano, E., Vodrazka, P., Kuner, R., Groene, H.J., Herr, I., Krammer, P.H. & Martin-Villalba, A. (2006) Control of neuronal branching by the death receptor CD95 (Fas/Apo-1). Cell death and differentiation, 13, 31–40.

